# Application of a GRF-GIF chimera enhances plant regeneration for genome editing in tomato

**DOI:** 10.1101/2024.12.20.628945

**Authors:** Gwen Swinnen, Eléonore Lizé, Miguel Loera Sánchez, Stéphanie Stolz, Sebastian Soyk

## Abstract

Genome editing has become a routine tool for functionally characterizing plant and animal genomes. However, stable genome editing in plants remains limited by the time- and labor-intensive process of generating transgenic plants, as well as by the efficient isolation of desired heritable edits. In this study, we evaluated the impact of the morphogenic regulator *GRF-GIF* on plant regeneration and genome editing outcomes in tomato. We demonstrate that expressing a tomato *GRF-GIF* chimera reliably accelerates the onset of shoot regeneration from callus tissue culture by approximately one month and nearly doubles the number of recovered transgenic plants. Consequently, the *GRF-GIF* chimera enables the recovery of a broader range of edited haplotypes and simplifies the isolation of mutants harboring heritable edits, but without markedly interfering with plant growth and development. Based on these findings, we outline strategies that employ basic or advanced diagnostic pipelines for efficient isolation of single and higher-order mutants in tomato. Our work represents a technical advantage for tomato transformation and genome editing, with potential applications across other Solanaceae species.

## Introduction

The application of genome editing using site-specific nucleases has become routine in many model plants and crops to study gene function. The most common approach to deliver clustered regularly interspaced short palindromic repeats-CRISPR associated protein (CRISPR-Cas) reagents to plant cells is the stable integration of a transgene that encodes for the site-specific Cas nuclease enzyme and guide RNAs (gRNAs) through *Agrobacterium*-mediated transformation (B. Li *et al*., 2024). However, the stable transformation of plants still presents a limiting factor for genome editing experiments, especially in species that require tissue culture transformation. Laborious and inefficient transformation protocols are a particular bottleneck for advanced editing applications such as base editing, prime editing, gene targeting, *cis*-regulatory editing, multiplex editing, and CRISPR screens, which require not only the generation of large cohorts of independent transgenic events but also an efficient and user-friendly approach to analyze and recover desired editing outcomes.

For many species, the rate-limiting step is not the transformation of plant cells *per se* but rather the regeneration of viable first-generation (T0) transgenic plants. Expression of morphogenic regulators has been shown to improve plant regeneration in species and genotypes that are recalcitrant to regeneration in tissue culture. A prominent example is the combined expression of BABYBOOM (BBM) and WUSCHEL (WUS) homologs in maize (*Z. mays*) and other monocots (Lowe *et al*., 2016). WUS is a homeobox transcription factor and key regulator of meristem identity (Zuo *et al*., 2002), while BBM is a member of the AP2/ERF transcription factor family that plays a crucial role during embryogenesis (Boutilier *et al*., 2002). Ectopic expression of BBM or WUS leads to the formation of somatic embryos and combined expression stimulates growth of embryogenic callus tissue (Boutilier *et al*., 2002; Lowe *et al*., 2016; Zuo *et al*., 2002). However, ectopic BBM and WUS expression has pleiotropic effects including severe changes in plant morphology and reduced fertility (Lowe *et al*., 2016). Overcoming these detrimental effects requires excision of *BBM* and *WUS* using the Cre/loxP system after stimulation of embryogenic callus formation or the use of tissue and developmental-stage specific promoters (Lowe *et al*., 2016).

More recently, constitutive expression of a chimeric protein composed of the GROWTH-REGULATING FACTOR 4 (GRF4) and its cofactor GRF-INTERACTING FACTOR 1 (GIF1) has been shown to improve regeneration in wheat, rice, and citrus without detrimental effects on plant growth and development (Debernardi *et al*., 2020). GRF4 belongs to a family of transcription factors that are deeply conserved across the plant kingdom and act as positive regulators of cell proliferation (Omidbakhshfard *et al*., 2015). GRF transcription factors interact with GIF co-activators that enable the formation of a complex with chromatin remodeling factors to regulate cellular growth and organ development (Debernardi *et al*., 2014; Nelissen *et al*., 2015). It was proposed that synthetic *GRF-GIF* chimeras enhance plant regeneration due to its capacity to drive cell proliferation across multiple organs and its role in regulating the transition of stem cells to transit-amplifying cells, which typically undergo rapid cell division to ensure the availability of sufficient cells before differentiation (Debernardi *et al*., 2020; Rodriguez *et al*., 2010, 2015).

Since the first report in wheat and citrus (Debernardi *et al*., 2020), *GRF-GIF* chimeras have been applied in diverse species to enhance plant regeneration without any documented pleiotropic effects (Bull *et al*., 2023; Feng *et al*., 2021; J. Li *et al*., 2024; Vandeputte *et al*., 2024; Zhang *et al*., 2021; Zhao *et al*., 2024). However, whether *GRF-GIF* chimeras enhance regeneration in Solanaceae species including tomato (*Solanum lycopersicum*) has not been tested before. Tomato is an important fruit vegetable crop and acts as a model system for both fundamental and translational research. Efficient protocols for tomato transformation (van Eck *et al*., 2019) and CRISPR-Cas genome editing (Brooks *et al*., 2014) have been reported but regeneration efficiency can vary greatly between protocols and experiments. The intense time and labor investment required for tomato transformation, especially compared with the model species Arabidopsis, therefore remains a barrier for many genetic and mechanistic studies, underlining the need to further improve the efficiency of transformation protocols.

In this study, we show that ectopic expression of a *S*. *lycopersicum GRF-GIF* (*SlGRF-GIF*) chimera enhances regeneration efficiency in tomato tissue culture by nearly two-fold while also reducing the time to obtain transgenic plants by approximately four weeks. Our results demonstrate that simultaneous delivery of the SlGRF-GIF complex together with genome editing reagents yields higher mutant haplotype diversity without markedly interfering with heritability or plant development.

## Results

### Identification of *GRF4* and *GIF1* homologs in tomato

To identify *GRF4* and *GIF1* homologs in tomato, we conducted a phylogenetic analysis with GRF and GIF proteins from Arabidopsis, grapevine (*V. vinifera*), citrus (*C. clementina*), rice (*O. sativa*), maize (*Z. mays*) and tomato. Consistent with previous studies in Arabidopsis, we found that tomato GIF proteins grouped into two subclades based on homology to GIF1 and GIF2/GIF3 (**Fig. S1a**) (Kim and Kende, 2004). The two tomato GIF proteins, which clustered closely to GIF1 from Arabidopsis, grapevine and citrus, were named *S. lycopersicum* GIF1a (SlGIFa) and SlGIF1b. Both *SlGIF1a* and *SlGIF1b* transcripts were primarily expressed in meristematic tissue with overall higher expression levels for *SlGIF1b* (**Fig. S1b**). From the 13 annotated tomato GRF (SlGRF) proteins we identified three with high sequence similarity to the GRF4 orthologs from grapevine and citrus, which we named SlGRF4a-c (**Fig. S1c**) (Debernardi *et al*., 2020). All three *SlGRF4* homologs were primarily expressed in meristematic tissue with similar expression patterns across different stages of meristem maturation (**Fig. S1d**). We constructed a chimeric SlGRF4c-SlGIF1a (SlGRF-GIF) protein with an alanine linker according to Debernardi and colleagues (**Fig. S1e**) (Debernardi *et al*., 2020), and combined the *SlGRF-GIF* chimera with a parsley *UBIQUITIN* promoter (pPcUbi) and the *Pisum sativum rbcS-3A* gene terminator (Pea3At) in an expression module for the widely used Golden Gate Modular Cloning (MoClo) system (Engler *et al*., 2014; Werner *et al*., 2012).

### *SlGRF-GIF* improves regeneration efficiency of tomato

To determine whether the *SlGRF-GIF* chimera enhances regeneration of tomato plants from tissue culture, we combined the *SlGRF-GIF* chimera with a kanamycin resistance marker (*NptII*) and the visual *RUBY* reporter (He *et al*., 2020) (**Fig. 1a**). *RUBY* is a synthetic open reading frame encoding three enzymes (CYP76AD1, DODA, and Glucosyltransferase) that catalyze subsequent steps in the synthesis of the red pigment betalain. We introduced the construct into tomato cotyledon explants using *Agrobacterium*-mediated transformation. In parallel, we transformed a construct without the *SlGRF-GIF* expression cassette as a control. To assess regeneration efficiency, we quantified the formation of shoots that developed on explants transformed with either the *SlGRF-GIF* or control constructs (**Fig. 1b** and **Fig. S2a-d**). We observed 1.8-fold more shoots on *SlGRF-GIF* explants (45 shoots) compared with the control (25 shoots) eight weeks after transformation (**Fig. 1b**), and consistently more shoots on *SlGRF-GIF* explants throughout the experiment. A similar increase in shoot regeneration from the *SlGRF-GIF* construct became apparent at the transfer to rooting medium. After ten weeks, we observed 1.8-fold more shoots at this step for the *SlGRF-GIF* construct (31 shoots) compared with the control (17 shoots) (**Fig. 1c**). Importantly, the *SlGRF-GIF* construct also yielded more rooted regenerants that acclimated on soil. Within the first twelve weeks (3 months), we obtained 8 and 3 rooted regenerants on soil for the *SlGRF-GIF* and the control construct, respectively (**Fig. 1d**). In total, we recovered 26 plants from the *SlGRF-GIF* construct and 14 plants from the control before we stopped the experiment after 18 weeks (4.5 months) (**Fig. 1e**). Based on these observations, we estimated regeneration efficiency as the number of regenerated plants per number of inoculated explants and found that regeneration efficiency with the *SlGRF-GIF* construct (13%) was 1.9-fold higher compared with the control (7%) (**Fig. 1e,f**).

**Fig. 1:**
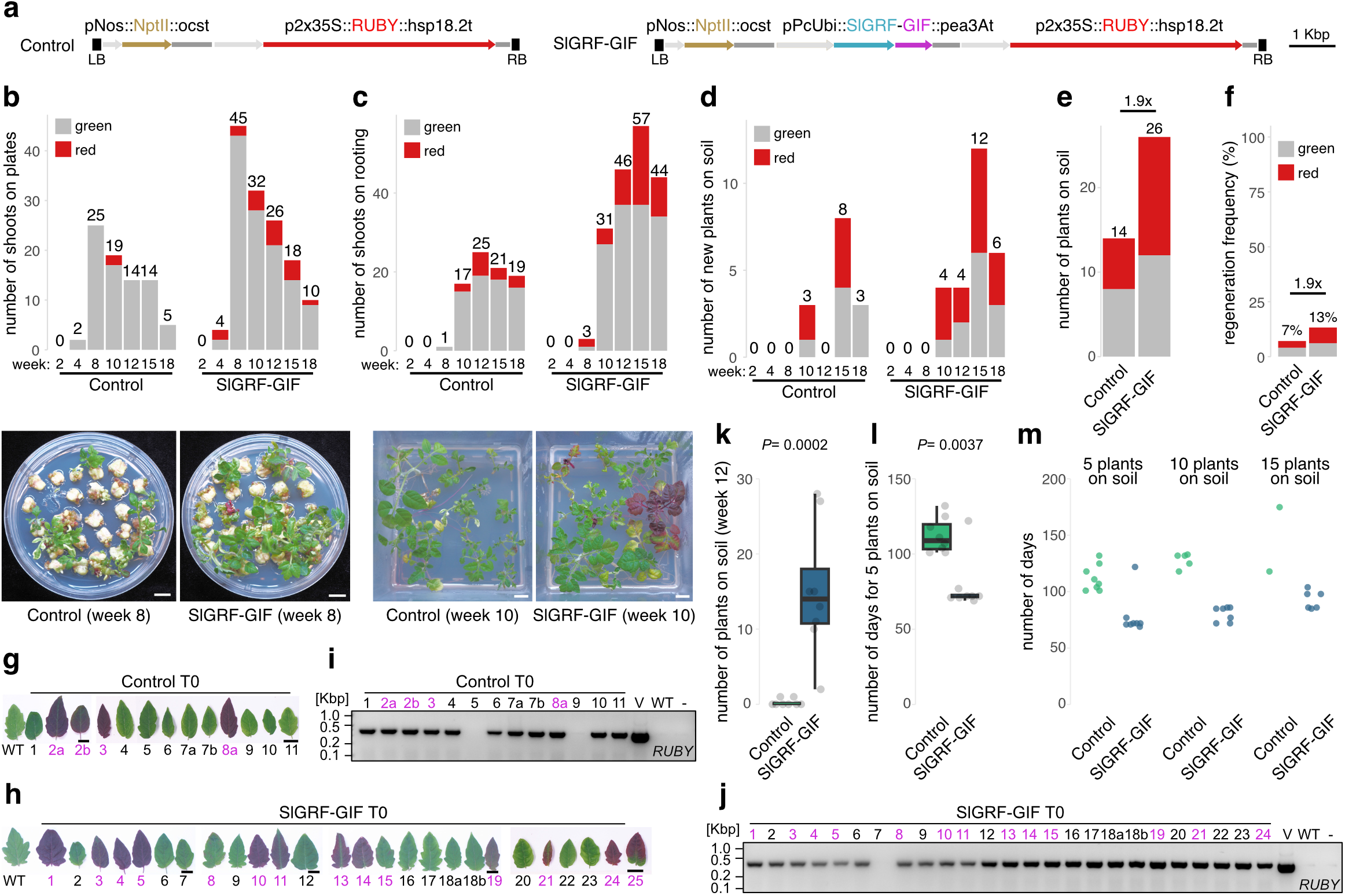
Assessing the effect of the *SlGRF-GIF* chimera on tomato plant regeneration in combination with the visible *RUBY* reporter. **a**, Configuration of the control and *SlGRF- GIF* constructs. LB, left border; RB, right border; *pNos*, *A. tumefaciens nopaline synthase* promoter; *NptII*, *E. coli neomycin phosphotransferase II* (Kanamycin resistance); *ocst*, *A. tumefaciens octopine synthase* terminator; *p2x35S*, double cauliflower mosaic virus 35S promoter; *RUBY*, synthetic *RUBY* reporter; *hsp18.2t*, *A. thaliana HSP18.2* terminator; *pPcUbi*, *P. crispum UBIQUITIN* promoter; *SlGRF-GIF*, *S. lycopersicum GRF4c-GIF1a* chimera; *pea3At*, *P. sativum rbcS-3A* terminator. **b**, Quantification of green and red shoots on regeneration medium per week. Images of representative plates during week 8 are shown. **c**, Quantification of green and red shoots on rooting medium per week. Images show representative boxes during week 10. **d**, Quantification of new green and red regenerants transferred to soil per week. **e**, Total number of green and red regenerants transferred to soil by the end of the experiment after 18 weeks. **f**, Regeneration efficiency for plants on soil (total number of plants by the number of explants at the start of the experiment) for the control and *SlGRF-GIF* experiments. **g-h**, Images of detached leaflets from individual regenerants from the control (g) and *SlGRF-GIF* (h) experiment illustrating the red pigmentation from *RUBY* activity. **i-j**, Detection of the *RUBY* transgene by PCR and agarose gels in control (i) and *SlGRF-GIF* (j) regenerants with vector (V), wild type (WT), and no DNA (-) control. Kbp, kilo base pairs. **k**, Number of regenerants on soil 12 weeks after transformation for 8 independent experiments with and without *SlGRF-GIF*. **l-m**, Number of days to obtain 5 (l-m), 10, and 15 (m) regenerants on soil for 8 independent experiments with and without *SlGRF-GIF*. Numbers above bars in (b-e) indicate the total number of regenerants (red and green) per week. Scale bars in (b-c, g-h) represent 1 cm. Magenta font in (g-j) indicates regenerants with red pigmentation. *P* values in (k,l) indicate results from a two-tailed, two sample Mann-Whitney test.

Surprisingly, only 53.8% of *SlGRF-GIF* regenerants (14/26) and 42.8 % of control regenerants (6/14) developed the red pigmentation that indicates *RUBY* activity (**Fig. 1e,g-h**). In addition, pigmentation intensity varied considerably between T0 individuals (**Fig. S2e-f**). We reasoned that regenerants without the red pigmentation may represent failed transformation events that escaped antibiotic selection and, therefore, tested for the genomic integration of the *RUBY* coding sequence by PCR (**Fig. 1i-j**). However, only two control (T0-5 and T0-9) and a single *SlGRF-GIF* transformant (T0-7) tested negative for *RUBY* while all other green plants carried the transgene. We concluded that the high fraction of transgenics without visible *RUBY* activity was likely due to silencing of the *RUBY* transgene. Importantly, overexpression of *SlGRF-GIF* did not obviously interfere with plant growth and development, and we obtained T1 seeds from nearly all plants except some with strong *RUBY* coloration. This initial experiment suggested that the *SlGRF-GIF* chimera can be used to increase regeneration frequency in tomato, and in turn, the number of confirmed transgenic events.

To determine whether the *SlGRF-GIF* chimera reliably enhances regeneration, we evaluated eight independent transformation experiments with and without *SlGRF-GIF* that we performed over the course of two and a half years. Within the first twelve weeks (3 months) after transformation, we obtained on average 0.25 and 15 regenerants on soil for the control and *SlGRF-GIF* transformations, respectively (**Fig. 1k**). Remarkably, transformations with *SlGRF- GIF* yielded the first five regenerated plants on average 34 days earlier than the controls (**Fig. 1l**), and a similar trend was observed for the first 10 and 15 plants (**Fig. 1m**). We also noted that fewer control transformations produced at least 10 or 15 plants compared with *SlGRF-GIF* transformations (**Fig. 1m**). These results demonstrate that *SlGRF-GIF* consistently enhances shoot regeneration and shortens the time to obtain regenerated plants on soil by approximately four weeks.

### *SlGRF-GIF* streamlines the production of CRISPR-Cas mutants

Next, we tested whether the combination of the *SlGRF-GIF* chimera and the *RUBY* reporter can simplify the generation of tomato mutants by CRISPR-Cas mutagenesis. We speculated that intensity of *RUBY* coloration could be used as a proxy for the expression of transgenes from the T-DNA, and thus indirectly for Cas9 expression and mutagenic activity. This could allow the selection of red transgenics during early stages of the transformation process to reduce the number of transformants for molecular analyses. We constructed a binary acceptor construct containing *SlGRF-GIF*, *RUBY*, the Cas9 nuclease, and an array of three gRNAs to target the florigen-encoding *SINGLE FLOWER TRUSS* (*SFT*) gene (Lifschitz *et al*., 2006) (**Fig. 2a**, **Fig. S3a**). Within 14 weeks (3.5 months), we recovered 13 regenerants on soil, however, four plants (31%) did not develop the red pigmentation although the transgene was confirmed by PCR (**Fig. 2b,c**). We also PCR-amplified the *SFT* target region for quick and easy detection of Cas9-induced DNA lesions on agarose gels (**Fig. 2d**). DNA lesions were clearly visible in most transgenics, but they were not reliably indicated by the red pigmentation from *RUBY* activity (**Fig. 2e**). In fact, Sanger sequencing of target amplicons revealed mutation rates of at least 75% in each transgenic plant (**Table S1**), and confirmed homozygous edits in two individuals (T0-6 and T0-10) (**Fig. S3b**). We also asked whether *RUBY* activity correlated with phenotypic changes expected from the Cas9-induced mutations and scored plants that developed *sft* mutant inflorescences, which are single-flowered or revert to vegetative growth (Lifschitz *et al*., 2006). In addition, we scored T0 plants that reverted from determinate to indeterminate shoot growth due reduced *SFT* activity (Krieger *et al*., 2010) (**Fig. S3c-d**). Supporting the high mutation rates detected by Sanger sequencing, we observed strong *sft* mutant phenotypes in 69-77 % of the T0 plants (**Fig. 2f,g** and **Fig. S3c-d**). However, the changes were not obviously associated with *RUBY* coloration. For example, the green *sft^CR^*-4 individual developed the characteristic *sft* single-flowered truss while the inflorescences of the red *sft^CR^*-5 individual were indistinguishable from the WT (**Fig. 2f**).

**Fig. 2:**
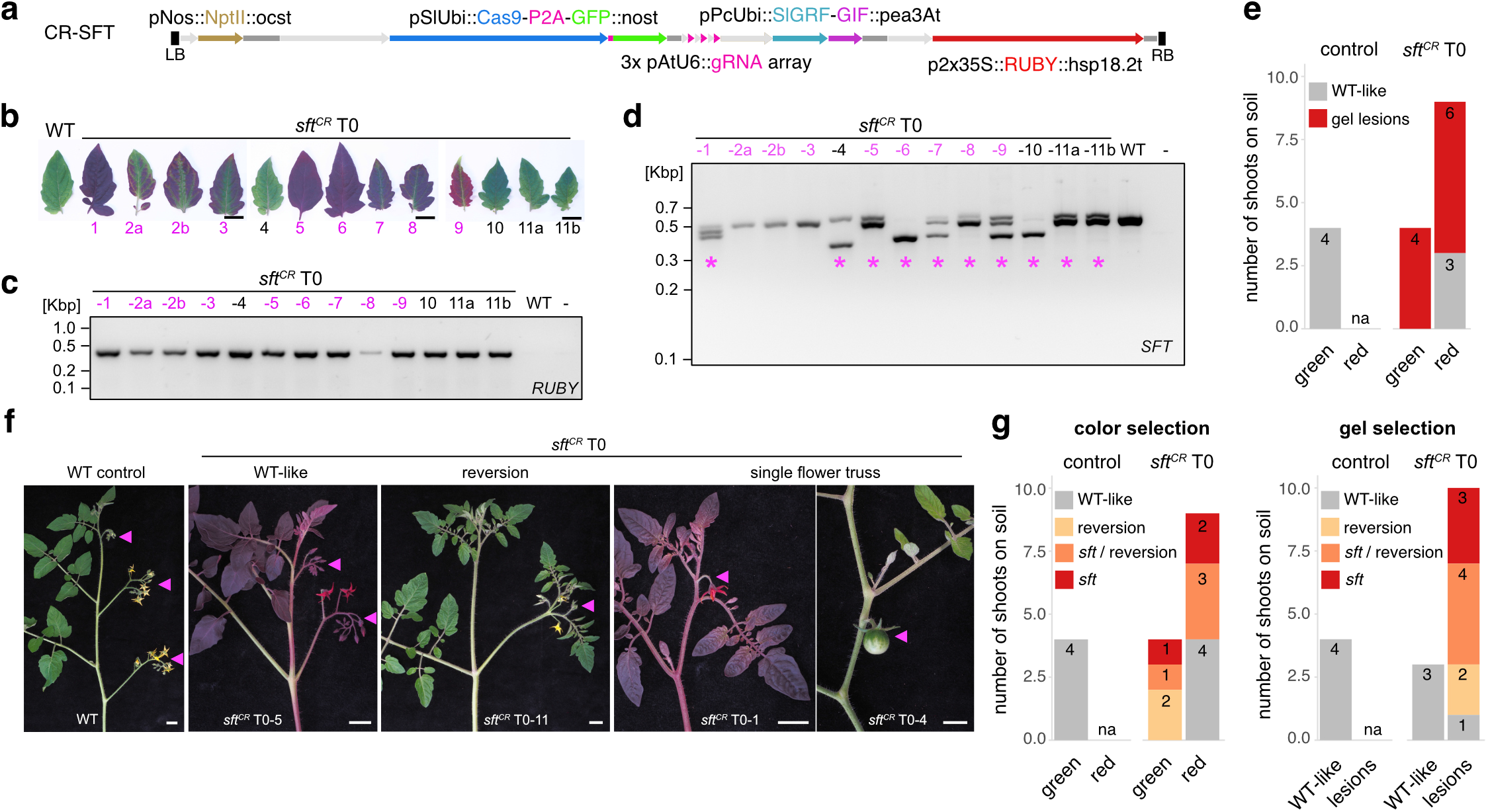
Application of the *SlGRF-GIF* chimera for CRISPR-Cas targeting of *SFT* in combination with the visible *RUBY* reporter. **a**, Construct configuration for targeting the *SFT* gene by CRISPR-Cas. LB, left border; RB, right border; *pNos*, *A. tumefaciens nopaline synthase* promoter; *NptII*, *E. coli neomycin phosphotransferase II*(Kanamycin resistance); *ocst*, *A. tumefaciens octopine synthase* terminator; *pSlUbi*, *S. lycopersicum UBIQUITIN* promoter; *Cas9-P2A-GFP*; *S. pyogenes cas9* nuclease linked to GFP by the self-cleaving peptide P2A; 3x pAtU6::gRNA; 3 gRNA expression cassettes driven by *A. thaliana* U6 promoter. pPcUbi, *P. crispum UBIQUITIN* promoter; *SlGRF-GIF*, *S. lycopersicum GRF4c-GIF1a* chimera; *pea3At*, *P. sativum rbcS-3A* terminator; *p2x35S*, double cauliflower mosaic virus 35S promoter; *RUBY*, synthetic *RUBY* reporter; *hsp18.2t*, *A. thaliana HSP18.2* terminator. **b**, Images of detached leaflets from the wild type (WT) and 13 *sft^CR^* regenerants. Note that *sft^CR^*-2a/b and *sft^CR^*-11a/b originate from the same callus. **c**, Detection of the *RUBY* transgene by PCR amplification and agarose gels in *sft^CR^* regenerants and the WT and no DNA (-) controls. **d**, Detection of Cas9- induced DNA lesions in *SFT* by PCR amplification and agarose gels in *sft^CR^*regenerants and the WT and no DNA (-) controls. Magenta asterisks mark samples with clearly visible DNA lesions. **e**, Quantification of the number of samples with detectable DNA lesions for the WT control and first-generation (T0) transgenic *sft^CR^* lines. Individuals were grouped by plant RUBY coloration. The number of individual plants per category is indicated. **f**, Images of representative shoots from the WT control and selected *sft^CR^* regenerants to illustrate observed inflorescence phenotypes (WT-like, reversion, and single flower truss). Magenta arrowheads mark inflorescences. Note that *sft* mutant phenotypes are observed on both red and green plants. **g**, Quantification of plants with mutant inflorescence phenotypes for the WT control and first- generation (T0) transgenic *sft^CR^* lines. Individuals were grouped by RUBY coloration (left) or the presence of clearly visible DNA-lesions on agarose gels (right). The number of individual plants per category is indicated. Scale bars in (b, f) represent 1 cm. Kbp in (c-d), kilo base pairs. Magenta font in (b-d) indicates regenerants with red pigmentation. na in (e, g), not applicable.

In conclusion, we could not exploit *RUBY* as a proxy for Cas9 activity or the frequency of Cas9-induced mutations, most likely due to silencing of the *RUBY* transgene. We also observed that strong *RUBY* coloration negatively affected plant growth and fertility, and most plants with strong red pigmentation failed to set seed. Such side effects from betalain overproduction have been reported in tomato (Wang *et al*., 2023), and it remains to be tested whether weaker or tissue-specific promoters allow to establish *RUBY* as reliable selection marker in tomato. Nevertheless, the SlGRF-GIF fusion protein did not obviously interfere with CRISPR-Cas activity as mutant phenotypes could be observed at high frequency in the first (T0) generation of transgenics.

To test whether enhanced plant regeneration from *SlGRF-GIF* could improve the isolation of higher-order mutants by CRISPR-Cas, we generated constructs with and without *SlGRF-GIF* to simultaneously target the MADS-box transcription factor genes *JOINTLESS2* (*J2*) and *ENHANCER OF J2* (*EJ2*) with two gRNAs per target gene (**Fig. 3a** and **Fig. S4a,d**). Individual mutation of *J2* and *EJ2* affect the development of fruit abscission zones and sepals, respectively, while simultaneous mutation of *J2* and *EJ2* causes strong inflorescence branching (Soyk et al., 2017). Eight weeks after transformation, we observed that explants transformed with the *SlGRF-GIF* construct produced 1.9-fold more shoots than the control explants (**Fig. 3b,c**). The *SlGRF-GIF* construct also extended the duration of shoot regeneration, which resulted in 2.2- fold more shoots (53 versus 24) by the end of the experiment after 15 weeks (**Fig. 3d**) and translated to shoot regeneration efficiencies of 24% and 8% for the *SlGRF-GIF* and the control experiment, respectively (**Fig. 3e**). The *SlGRF-GIF* experiment also yielded more shoots for the transfer to rooting medium during weeks 8-9 (**Fig. 3f,g**), which increased the total number of shoots on rooting medium from 19 to 40 (**Fig. 3h**) and enhanced regeneration efficiency at this step from 8% to 16% compared with the control (**Fig. 3i**). However, only 5 and 13 shoots rooted successfully for the control and *SlGRF-GIF* transformations, respectively, corresponding to a regeneration efficiency of 2.0% and 5.3% at the plant level (**Fig. 3k,l**). We reasoned that this relatively low regeneration rate may be due to the fact that *J2* and *EJ2* are regulators of meristem development. Interestingly, all 13 rooted *SlGRF-GIF* regenerants developed branched (compound) inflorescences that are characteristic of *j2ej2* double mutants whereas only 40% (2/5) of the control regenerants displayed the double mutant phenotype (**Fig. 3j**,**k**).

**Fig. 3:**
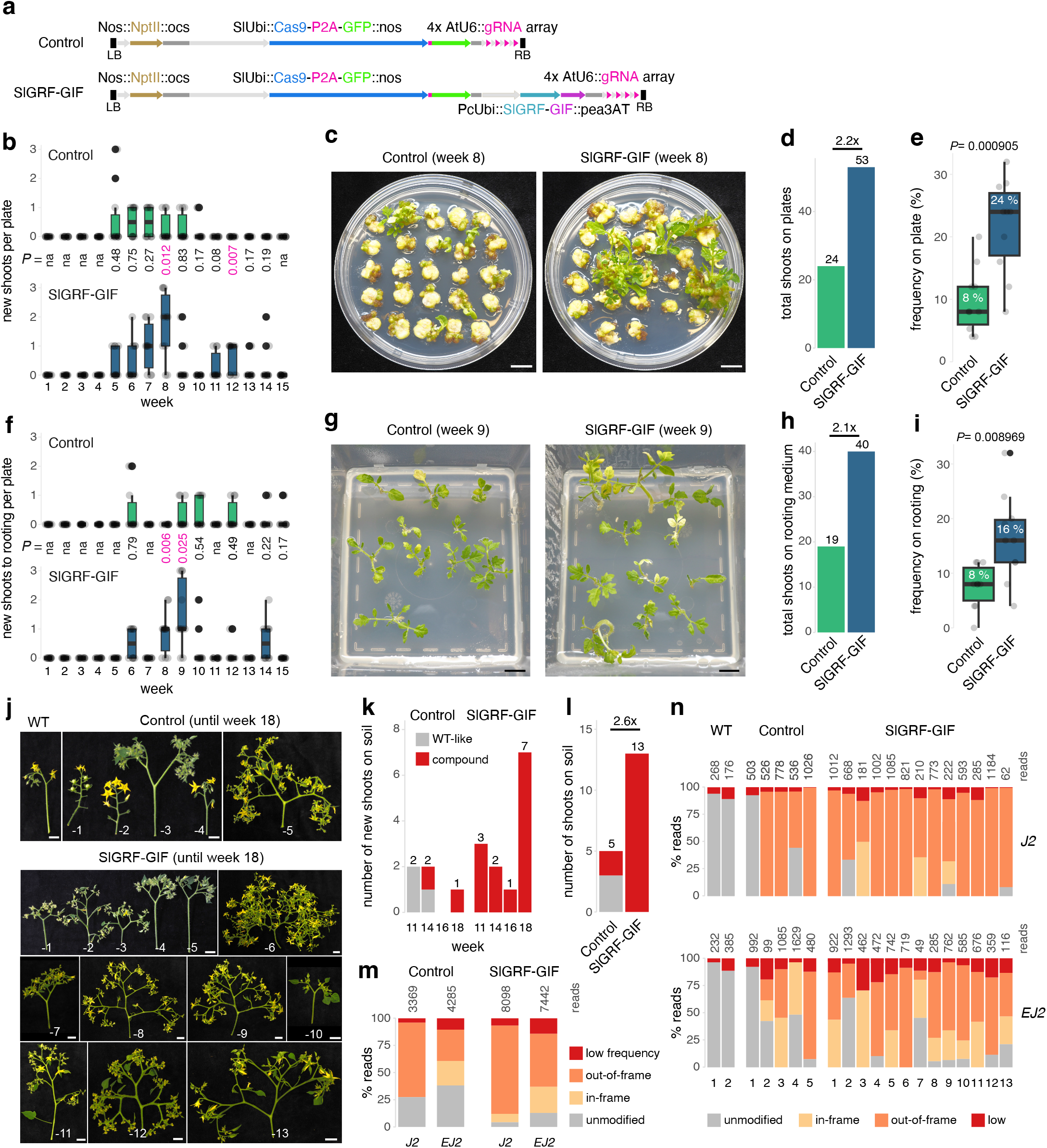
CRISPR-Cas multiplex targeting of *J2* and *EJ2* with and without the *SlGRF-GIF* chimera. **a,** Configuration of the control and *SlGRF-GIF* constructs for targeting the *J2* and *EJ2* genes by CRISPR-Cas. LB, left border; RB, right border; *pNos*, *A. tumefaciens nopaline synthase* promoter; *NptII*, *E. coli neomycin phosphotransferase II* (Kanamycin resistance); *ocst*, *A. tumefaciens octopine synthase* terminator; *pSlUbi*, *S. lycopersicum UBIQUITIN* promoter; *Cas9-P2A-GFP*; *S. pyogenes cas9* nuclease linked to GFP by the self-cleaving peptide P2A; 3x pAtU6::gRNA; 4 gRNA expression cassettes driven by *A. thaliana* U6 promoter. *pPcUbi*, *P. crispum UBIQUITIN* promoter; *SlGRF-GIF*, *S. lycopersicum GRF4c-GIF1a* chimera; *pea3At*, *P. sativum rbcS-3A* terminator. **b**, Quantification of shoots that emerged on regeneration medium per week. Each datapoint indicates the number of shoots that emerged on an individual plate (n=10). **c**, Images show representative plates with regeneration medium for the control and *SlGRF-GIF* experiment during week 8. **d**, Quantification of shoots that emerged on regeneration medium until the end of the experiments. **e**, Regeneration frequency for shoots on regeneration medium (total number of shoots by the number of explants at the start of the experiment). **f**, Quantification of new shoots transferred to rooting medium per week. Each datapoint represents the number of new shoots transferred from a single plate (n=10). **g**, Images show representative rooting medium boxes for the control and *SlGRF-GIF* experiment during week 9. **h**, Quantification of regenerated shoots transferred to rooting medium until the end of the experiments. **i**, Regeneration frequency for shoots on rooting medium (total number of shoots by the number of explants at the start of the experiment). **j**, Images of representative inflorescences from *j2^CR^ej2^CR^* first generation (T0) transgenics for the control and *SlGRF-GIF* experiments transferred to soil until the end of the experiment (week 18). **k**, Quantification of new shoots transferred to soil per week. Differences in inflorescence development (WT-like or compound inflorescence) are indicated. **l**, The total number of regenerated shoots by the end of the experiment (week 18) as in (k). **m**, Mutation frequency at *J2* and *EJ2* in the pools of control and *SlGRF-GIF* T0 transgenics determined by amplicon sequencing. **n**, Mutation frequencies for *J2* and *EJ2* in individual control and *SlGRF-GIF* T0 transgenics determined by amplicon sequencing. Numbers on top of the bars in (m,n) indicate the number of aligned reads. Scale bars in (c,g,j) represent 1 cm. *P* values in (b, e, f, i) indicate results from a two-tailed, two sample t-test.

To identify T0 individuals that contain loss-of-function haplotypes at high frequency, we genotyped the gRNA target sites in *J2* and *EJ2* using amplicon sequencing and called haplotypes with the SMAP-package (Develtere *et al*., 2023; Schaumont *et al*., 2022). As expected for chimeric T0 plants, we detected genotypes that ranged from genetic mosaics to biallelic and homozygous edits for both *J2* and *EJ2* (**Fig. S4c,f** and **Table S2**). Interestingly, the *SlGRF-GIF* experiment yielded a larger number of unique haplotypes for both targets (**Fig. S4c,f**). For *J2*, we detected three unique mutant haplotypes in the control and 18 in the *SlGRF- GIF* regenerants. Similarly, for *EJ2*, we identified six unique haplotypes in the control and 20 in the *SlGRF-GIF* regenerants. However, the most abundant haplotypes for both genes were identical between experiments. For *J2*, this haplotype contained a 95 bp deletion, while the most abundant *EJ2* haplotype was the unmodified wild-type (WT) sequence (**Fig. S4c,f**). Overall, we detected a higher mutation frequency for *J2* than *EJ2*, but mutation frequencies did not significantly differ between the control and *SlGRF-GIF* experiments (**Fig. 3m** and **Table S3**). These results show that the *SlGRF-GIF* chimera does not interfere with genome editing efficiency but indirectly increases the number of mutant haplotypes by yielding a larger number of transgenic events.

To fix mutant haplotypes in the next generation (T1), we selected T0 individuals with high mutation frequencies. Among the *SlGRF-GIF* regenerants, T0-5 contained a homozygous (>85% haplotype frequency) out-of-frame edit for *J2* (*j2^h06^*) and two high-frequency *EJ2* haplotypes (*ej2^h04^* and *ej2^h15^*) (**Fig. 3n**, **Fig. S4c,f**, and **Table S2**). Furthermore, the *SlGRF-GIF* T0-6 displayed homozygous out-of-frame edits for both *J2* (*j2^h06^*) and *EJ2* (e*j2^h14^*) (**Table S2**). In the control experiment, we did not identify any T0s with homozygous out-of-frame mutations in both target genes. However, T0-3 was homozygous for *J2* (*j2^h06^*) and biallelic for *EJ2* (*ej2^h07^* and *ej2^h19^*), while T0-5 harbored biallelic out-of-frame edits for *J2* (*j2^h06^* and *j2^h08^*) and a near-homozygous out-of-frame edit (80.4% frequency) for *EJ2* (*ej2^h13^*) (**Fig. 3n**, **Fig. S4c,f**, and **Table S2**). Unfortunately, the control T0-5 plant did not set any fruits and seeds, most likely due to strong inflorescence branching.

We continued with three independent T1 families derived from the control (T0-3) and *SlGRF- GIF* (T0-5 and T0-6) regenerants, which showed high mutation frequencies for both target genes (**Fig. 3n** and **Fig. S5a**). All three T1 families segregated the T-DNA transgene, which suggested single insertion events although the frequency of transgenic individuals was lower than the expected 75% (**Fig. S5d**). We transplanted all non-transgenic plants and observed strong inflorescence branching on each individual (**Fig. S5e**), indicating homozygous or biallelic loss-of-function mutations in both target genes. To identify the transmitted haplotypes, we Sanger-sequenced selected target amplicons from non-transgenic plants (**Fig. S5d**). In the control T1-3 family, only a single *J2* haplotype (*j2^h06^*) was recovered, which had already been detected at a 96% frequency in the T0-3 parent (**Table S2**). For *EJ2*, we recovered two haplotypes (*ej2^h07^*and *ej2^h19^*) that had frequencies of 44.6% and 45.6% in the T0-3 parent. In the *SlGRF-GIF* T1-5 family, we again recovered the *j2^h06^* haplotype (T0-5 frequency of 97.1%), but we also isolated a large structural rearrangement (*j2^h22^*) that was not detected by amplicon sequencing in the T0-5 parent (**Fig. S5b**). For *EJ2*, only a low frequency (12.1%) haplotype (*ej2^h21^*) was recovered. In *SlGRF-GIF* T1-6, we recovered the *j2^h06^* and *ej2^h14^* haplotypes, which were detected at high frequencies (97.2% and 90.2%) in the T0-6 parent. However, we also identified a 16 bp/3 bp deletion in *J2* (*j2^h21^*) and a 56 bp deletion in *EJ2* (*ej2^h25^*) that were not detected in the T0-6 parent (**Fig. S5b-d**). These haplotypes were identified in non-transgenic T1 plants and were already visible on agarose gels in T0 parents (**Fig. S4b,e**), suggesting that they were already induced in the T0 generation.

We concluded that genotyping T0 individuals by amplicon sequencing can improve the selection of haplotypes that will be fixed in the T1 generation, but mutant haplotypes may be overlooked in the T0 generation if the experimental design is not customized to amplicon sequencing. Taken together, these results demonstrate that the *SlGRF-GIF* chimera facilitates CRISPR-Cas mutagenesis in tomato without compromising editing efficiency or heritability. In addition, the enhanced regeneration efficiency allows for greater haplotype diversity and easier selection of higher-order mutants in the T0 generation.

## Discussion

Despite continuous improvements of site-specific Cas nuclease variants and development of sophisticated genome editing applications, the rate-limiting step for CRISPR-Cas experiments in many plant species remains the ability to recover transgenic plants with heritable genome edits. The expression of morphogenic regulators during tissue culture has been shown to boost plant regeneration (Debernardi *et al*., 2020; Kong *et al*., 2020; Lowe *et al*., 2016; Maher *et al*., 2020). In this study, we evaluated the emerging morphogenic regulator GRF-GIF complex to enhance regeneration and streamline CRISPR-Cas experiments in tomato, a widely used model system for fundamental and translational research. Expression of a tomato *SlGRF-GIF* chimera significantly enhanced regeneration of transgenic tomato plants from cotyledon tissue culture. The chimeric protein reliably increased the efficiency of shoot regeneration by nearly two-fold and shortened the time to obtain transgenic plants by approximately four weeks, thereby reducing the time and labor investment for tissue culture. These findings support previous reports on the application of *GRF-GIF* chimeras in other species (Bull *et al*., 2023; Debernardi *et al*., 2020; Feng *et al*., 2021; J. Li *et al*., 2024; Vandeputte *et al*., 2024; Zhang *et al*., 2021; Zhao *et al*., 2024).

Importantly, ectopic *SlGRF-GIF* expression in tomato did not cause obvious detrimental effects on plant development and fertility, although we occasionally observed slightly elongated fruits. Such subtle pleiotropic effects are not expected to impact genome editing experiments that involve transgene segregation and selection of non-transgenic plants in subsequent generations. Still, pleiotropic effects could be minimized by inducible or tissue-specific expression of *SlGRF-GIF* during early stages of plant regeneration. When generating stable transgenic lines, e.g., for genomic complementation or reporter constructs, co- transformation of *SlGRF-GIF* on a separate construct as recently shown in lettuce is a possibility (Bull *et al*., 2023). Finally, SlGRF-GIF activity could be further increased by interfering with the post-transcriptional regulation by miRNA396. *GRF-GIF* variants with silent mutations in the miR396 target site increased the frequency of transgenic events in citrus and lettuce (Bull *et al*., 2023; Debernardi *et al*., 2020). However, miR396 resistant *GRF-GIF* variants caused pleiotropic effects with many transgenic events producing large callus tissue that was unable to regenerate shoots.

Although the first protocols for tomato transformation have been established several decades ago (Van Eck *et al*., 2019; McCormick *et al*., 1986; Sun *et al*., 2006), we see an advantage in using the *SlGRF-GIF* chimera for tomato genome editing mainly in three ways: First, the recovery of the transgenic plants is accelerated, which allows earlier verification of editing outcomes and characterization of mutant lines, or adjusting the experimental design if necessary. Second, the production of larger populations of transgenic plants is simplified, which becomes especially important when applying precision editing tools with low efficiency, editing *cis*-regulatory regions to obtain very specific alleles, employing multiplex editing, or performing CRISPR screening. Third, the production of transgenic plants becomes more robust and reproducible, which enables research teams that are not specialized in plant transformation to reliably produce sufficient numbers of transgenic events in a timely manner.

Designing genome editing experiments to generate stable mutant lines requires careful consideration of the number of target genes, the genetic interactions to be explored (such as single or higher-order mutants), and the desired number of independent alleles (**Fig. S6**). These parameters determine the number of independent transformants to be generated (Van Huffel *et al*., 2022), which will inform both the genotyping and gRNA design. In low-throughput experiments, such as producing single loss-of-function mutants, the induction of large lesions allows low-input genotyping by PCR and agarose gels. When multiplex editing or precise small changes are desired, such as base edits or the disruption of *cis*-regulatory elements, next-generation sequencing-based genotyping becomes the method of choice. Here, rapid identification of haplotype sequences and frequencies in the T0 generation provides early insights into the overall editing outcome. In addition, it allows the selection of individuals with desired haplotype sequences at high frequency, which can be fixed in the next generation. In order to maximize the efficiency and accuracy of genotyping by amplicon sequencing, computational tools such as SMAP design (Develtere et al., 2023) for simultaneous gRNA and amplicon design are highly recommended.

Overall, we present a technical advantage for plant transformation and genome editing in tomato. Our findings add to the growing literature that *GRF-GIF* chimeras can be applied in diverse dicot species, including Solanaceae, to enhance plant regeneration.

## Materials and Methods

### Plant material and growth conditions

Seeds of *S. lycopersicum* cv. Sweet-100 (S100) double-determinate were from our own stocks (Alonge *et al*., 2022). Seeds were germinated on soil in 96-cell plastic trays. Plants were grown under long-day conditions (16-h light/ 8-h dark) in a greenhouse supplemented with artificial light from LED panels (∼200 umol m^-2^s^-1^), constant temperature (25°C) and relative humidity (50-60%). Plants were grown in 5L pots (2 per pot) under drip irrigation and standard fertilizer regimes. Tomato plants were pruned to the primary and one (proximal) axillary shoot. Phenotypic data was collected from T0 and T1 generation plants as indicated in the figures. Statistical analyses of phenotyping data were conducted in R.

### Molecular cloning

Vectors for plant transformation and CRISPR-Cas were assembled using the MoClo Golden Gate cloning system as previously described (Alonge *et al*., 2022; Engler *et al*., 2014; Werner *et al*., 2012). DNA fragments for cloning were amplified with KOD One PCR Master Mix (Sigma-Aldrich). The *pPcUbi* promoter was amplified from pDePPE (Perroud *et al*., 2022) with primers P500+P501 and cloned into pICH41295 to obtain pLL011. *SlGRF4* (Solyc12g096070) and *SlGIF1a* (Solyc04g009820) were amplified from young inflorescence cDNA (accession Sweet-100) using primers P1260+P1261 and P1262+P1263, respectively. Gel-purified amplicons were fused in a second PCR step using primers P1260+P1263 and cloned into pICH41308 to obtain pLL033. *Pea3At* was amplified using primers P502+P503 and pDePPE as template, and cloned into pICH41276 to obtain pLL012. pLL011, pLL033, and pLL012 were recombined into pICH47751 to obtain pLL034. *RUBY* was amplified from *35S::RUBY* (Wang *et al*., 2023)(addgene #160908) with primers P1658+P1659 and cloned into pICH41308 to obtain pSS135. The *AtHSP18.2* terminator was amplified from *35S::RUBY* with primers P1660+P1661 and cloned into pICH41276 to obtain pSS136. pICH51288 (2X35S), pSS135, and pSS136 were recombined into pICH47761 and pICH47791 to obtain pSS139 and pEL010, respectively. pICSL11024 (addgene #51144, Nos::NptII::ocs), pICH54022, pICH54033, pSS139, and pICH41780 were recombined into pICSL4723-P1 to obtain pEL013. pICSL11024, pICH54022, pLL034, pSS139, and pICH41780 were recombined into pICSL4723-P1 to obtain pEL014. For CRISPR-Cas9, a new L1 part (*SlUbi::SpCas9-P2A- GFP::nos* part; SS082) and L2 acceptor (*pSlUbi:Cas9-SlGRF-GIF-LacZ*; pEL001) was cloned. The *SlUbi* promoter (2075 bp upstream of Solyc07g064130) was amplified with primers P126+P150 from gDNA (accession Sweet-100) and cloned into pICH41295 to obtain pSS051. pSS051, pSS058 (pAGM1287-SpCas9) (Glaus *et al*., 2024), SS065 (pAGM1301_P2A-GFP) (Glaus *et al*., 2024), and pICH41421 were recombined into pICH47742 to obtain pSS082. pICSL11024, pSS082, pLL034, and pICH49277 (lacZ) were recombined into pICSL4723-P1 (addgene #86173) to obtain pEL001. SFT-sgRNA-1, SFT-sgRNA-2, SFT-sgRNA-3 were cloned into pICH47761, pICH47772, pICH47781, respectively. Three L1 gRNA-constructs were recombined with pEL010 and plCH50866 into pEL001 to obtain pEL011. J2-sgRNA-1, J2-sgRNA-2, EJ2-sgRNA-1, EJ2-sgRNA-2 were cloned into pICH47761, pICH47772, pICH47781, pICH47791, respectively. The four L1 gRNA-constructs were recombined with plCH50866 into pEL001 to obtain pEL006. The four L1 gRNA-constructs were recombined with pICSL11024, pSS082, and pICH41822 into pICSL4723-P1 to obtain pEL005. All primer and gRNA sequences are listed in **Table S4** and **Table S5**, respectively. Vectors used in this study are listed in **Table S6**.

### Plant transformation

Binary vectors were transformed into tomato by *Agrobacterium*-mediated transformation according to Gupta and Van Eck (Gupta and Van Eck, 2016) with minor modifications. Briefly, seeds were sterilized by a series of incubation for 10 min in 70% ethanol, 5 min in sterile H2O, 15 min in 1.3% bleach, and rinsed four times with sterile water before sowing on MS medium (4.4 g/L MS salts, 1.5 % sucrose, 0.8 % agar, pH 5.9) in Sterivent containers (Duchefa S1686.0480). After sowing, seeds were kept for 72 h in darkness at 28°C before transfer to a Percival growth chamber with white LEDs (16 h light/8 h dark, ∼50 μmol m^−2^ s^−1^, 25°C, 50% humidity). Cotyledons were excised 7–8 days after sowing, cut in half, and incubated on 2Z- medium at 28°C in the dark for 24 h before transformation. *A. tumefaciens* (AGL-1) were grown in LB medium supplemented with antibiotics and washed in MS-0.2% medium (4.4 g/L MS salts, 2% sucrose, 100 mg/L myo-inositol, 0.4 mg/L thiamine, 2 mg/L acetosyringone, pH5.8). Explants were co-cultivated with *A. tumefaciens* on 2Z- medium supplemented with 100 μg/L IAA for 48 h at 28°C in the dark and transferred to 2Z selection medium (supplemented with 150 mg/L kanamycin). Explants were cultivated in a Percival growth chamber under white LEDs (16h light/8h dark, ∼50 μmol m^−2^ s^−1^, 25°C, 50% humidity) transferred every two weeks to fresh 2Z selection medium until shoot regeneration. Shoots were excised and transferred to selective rooting medium (supplemented with 150 mg/L kanamycin) in Sterivent containers. Well-rooted shoots were transplanted to soil and acclimated in a Percival growth chamber under white LEDs (16 h light /8 h dark, ∼50 μmol m^−2^ s^−1^, 25°C, 50% humidity) before transfer to the greenhouse.

### Genotyping

Genomic DNA was prepared from homogenized leaf tissue using extraction buffer (pH 9.5) containing 0.1 M of tris(hydroxymethyl)aminomethane–hydrochloride (Tris–HCl), 0.25 M of potassium chloride (KCl), and 0.01 M of ethylenediaminetetraacetic acid (EDTA). This mixture was incubated at 95°C for 10 min and subsequently cooled at 4°C for 5 min. After addition of 3% (w/v) BSA, collected supernatant was used as a template for PCR. All primers used for genotyping are listed in **Table S4**.

For low-throughput genotyping of selected T0 regenerants, target regions were amplified with gene-specific primers (**Table S4**) and PCR amplicons were purified using ExoSAP-IT (Thermo Fisher Scientific). After Sanger sequencing the purified PCR amplicons, quantitative sequence trace data were decomposed using Inference of CRISPR Editing (ICE) CRISPR Analysis Tool (https://ice.synthego.com/#/).

### Amplicon sequencing

High throughput genotyping of T0 regenerants was performed using amplicon deep sequencing as previously described (Liu *et al*., 2021) with minor modifications. Briefly, gRNA target regions were amplified using gene-specific primers (**Table S4**) with common adapter sequences. Gene-specific amplicons were diluted ten-fold in H2O before they were used as template for the addition of sample-specific indexes using adapter primers with unique barcodes as described before (Liu *et al*., 2021). Equal volumes of each indexed amplicon reaction were pooled and subsequently gel-purified using the Monarch DNA Gel Extraction Kit (New England Biolabs). A single Illumina sequencing library was prepared using the xGen DNA Library Prep MC Kit (Integrated DNA Technologies) and sequenced on the MiSeq System (Illumina) at the Genome Technologies Facility (GTF) at UNIL.

Raw reads were trimmed using Trimmomatic (v0.39; PE LEADING:3 TRAILING:3 SLIDINGWINDOW:4:15 MINLEN:36) (Bolger *et al*., 2014; Hannon, 2010) with the TruSeq2 Illumina adapter sequences (ILLUMINACLIP: TruSeq2-PE.fa:2:30:10:1:FALSE). Trimmed reads were split by barcode using fastx_barcode_splitter.pl from FASTX-Toolkit (v0.0.14) (Hannon, 2010). Mate pairing from R1 and R2 FASTQs was done using a custom awk script that pairs forward and reverse barcode reads that share sequence identifiers. The sequences of custom barcodes and linkers were removed by trimming the first 27 bp of each read with a custom awk script. The resulting FASTQ files were named according to their barcode combination. The R1 and R2 barcode-sorted reads were merged using PEAR (Zhang *et al*., 2014) (v0.9.6).

Barcode-sorted reads were mapped to the genomic sequence of their corresponding gene target using HISAT2 (v2.2.1) (Kim *et al*., 2019). The reference genomic sequence of each gene was extracted from the reference genome S100 (v2.0) (Alonge *et al*., 2022) in the forward direction and started at the transcriptional start site (TSS). The resulting BAM files were sorted and indexed using SAMtools (v.1.17) (Li *et al*., 2009). Haplotype calling was performed separately for each amplicon using SMAP haplotype-window (v5.0.1) (Develtere *et al*., 2023; Schaumont *et al*., 2022) retaining haplotypes with a minimum frequency of 5%. Amplicon borders were manually assigned depending on the location of the forward and reverse primers. Border start and end positions were scaled so that position 1 corresponds to the TSS of the target gene. Haplotype read counts and haplotype frequencies were extracted from the resulting SMAP haplotype-window read count and frequency tables.

To obtain insertion and deletion (INDEL) counts, each haplotype sequence was separately aligned to the reference sequence of its corresponding amplicon using MAFFT (v7.505) (Katoh, 2002) with the *adjustdirection* flag. The number of insertions was the count of gaps introduced in the reference sequence in the resulting alignment; correspondingly, the number of deletions was the count of gaps in the haplotype sequence. Haplotypes were categorized in “unmodified” (no insertions or deletions), “in-frame” (total INDEL count being a multiple of three or zero), “out-of-frame” (total INDEL count not being a multiple of three nor zero), and “low frequency" (haplotypes with per plant frequency of <5%). Low frequency haplotypes were not considered for further analysis. Average read percentages per plant and haplotype category combination were calculated separately for control and *SlGRF-GIF* regenerants. They were calculated by taking the sum of read percentages for each haplotype category per plant divided by the number of plants of the corresponding group (control and *SlGRF-GIF*). Wild-type control plants were not considered for the calculation of average read percentages. A multiple sequence alignment (MSA) that included a) the reference genomic sequence and b) the unmodified, in-frame and out-of-frame haplotype sequences was produced for each amplicon using MAFFT as described above. MSAs were manually edited to optimize visualization. Summary statistics and visualizations were generated using RStudio (v2023.06.1+524) with R (v4.3.1) (R Core Team, 2021; RStudio Team, 2020).

### Phylogenetic analysis

GRF protein sequences were retrieved from iTAK (itak.feilab.net/). GIF protein sequences were retrieved from Plaza (https://bioinformatics.psb.ugent.be/plaza/) using Arabidopsis GIF1 (AT5G28640) as query. Protein sequences were aligned with MAFFT (v7.481) (Katoh, 2002) using default parameters. Maximum likelihood phylogenetic trees were constructed with IQ- Tree (v2.2.0.5; parameters -m MFP -bb 1000 -bnni -redo) (Minh *et al*., 2020) and displayed in FigTree (v1.4.4; http://tree.bio.ed.ac.uk/software/figtree/).

## Acknowledgment

We thank all members of the Soyk lab for helpful discussions; L. Lebeigle and L. Currat for technical support with molecular cloning and plant transformation; B. Tissot, L. Nerny, L. Keel, and T. Stupp for support with plant care; J. Marquis and J. Weber for support with sequencing; This work was supported by the University of Lausanne, the European Research Council (ERC) under the European Union’s Horizon 2020 research and innovation programme (ERC Starting Grant “EPICROP” Grant No. 802008) to S.So. and a Swiss National Science Foundation (SNSF) Project Grant (Grant No. 310030_212218) to S.So.

## Author contributions

G.S. and S.So. conceived the project and designed and planned experiments. G.S., E.L., S.St., and S.So. performed experiments and collected data. G.S., E.L., M.L.S., S.St., and S.So. analysed data. S. So. acquired project funding. G.S. and S. So. wrote the first draft of the manuscript. All authors read, edited, and approved the manuscript.

## Data availability

Raw Illumina sequencing data will be made available on NCBI SRA under the BioProject PRJNA1200640 at the time of publication. Seeds and the SlGRF-GIF construct are available on request from S. Soyk.

## Supplementary Figure Legends

**Fig. S1:**
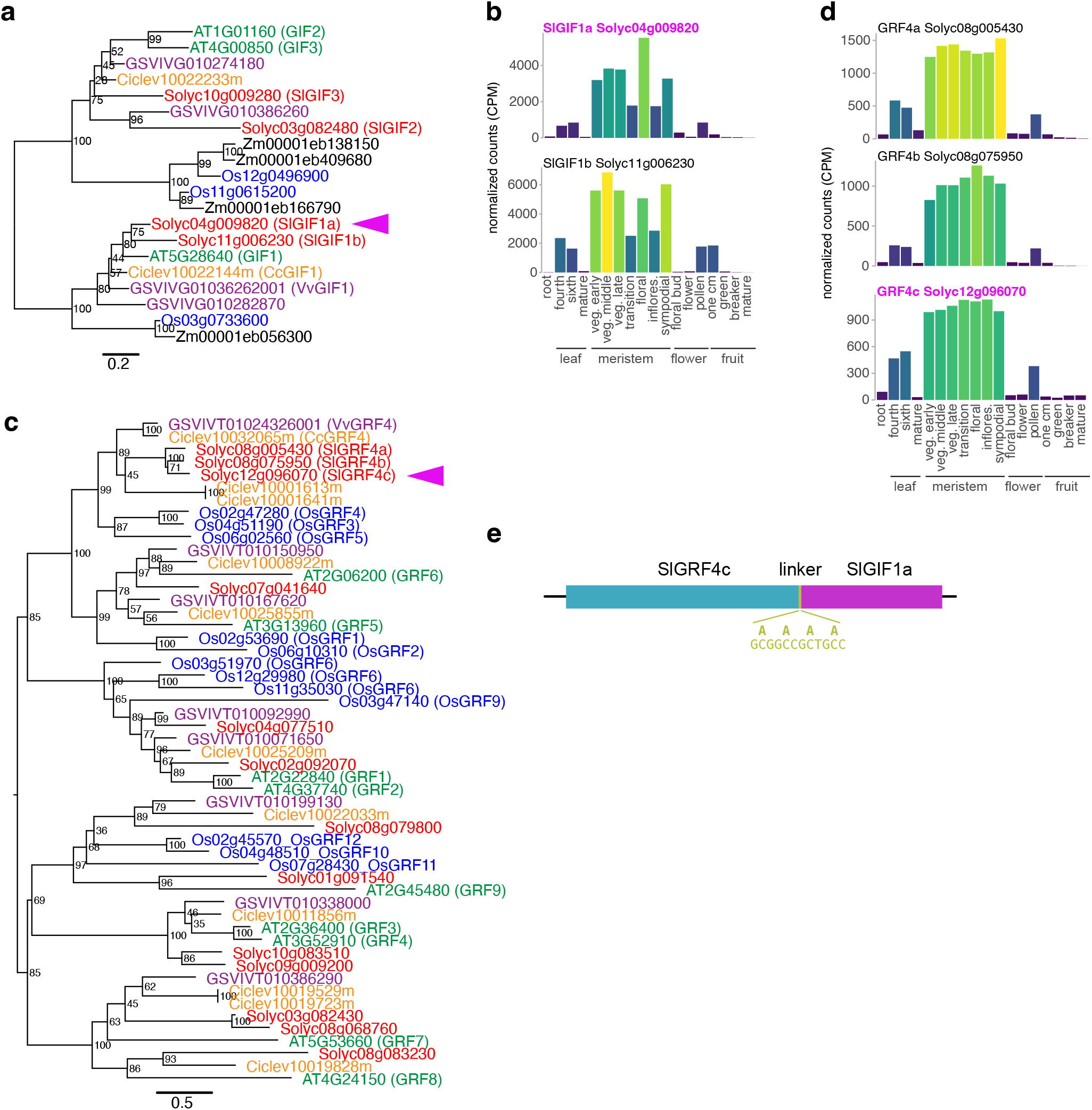
Identification of GRF4 and GIF1 homologs in tomato. **a**, **c**, Maximum likelihood phylogenetic tree for GIF (a) and GRF (b) protein homologs from *A. thaliana* (green), grape vine (*V. vinifera*, purple), citrus (*C. clementina*, orange), rice (*O. sativa*, blue), maize (*Z. mays*, black, only in GIF tree) and tomato (*S. lycopersicum*, red). Numbers at nodes indicate bootstrap values from 1000 replicates, and scale bars indicate the average number of substitutions per site. Tomato GIF and GRF homologs were named based on homology to Arabidopsis and *Vitis* proteins, respectively. **b-d**, Gene expression (RNA-seq) data for *SlGIF1* and *SlGRF4* genes in different tissues and developmental stages. CPM, counts per million. veg., vegetative. **e**, Schematic of SlGRF-GIF chimeric protein consisting of SlGRF4c and SlGIF1a linked by a polyalanine peptide.

**Fig. S2:**
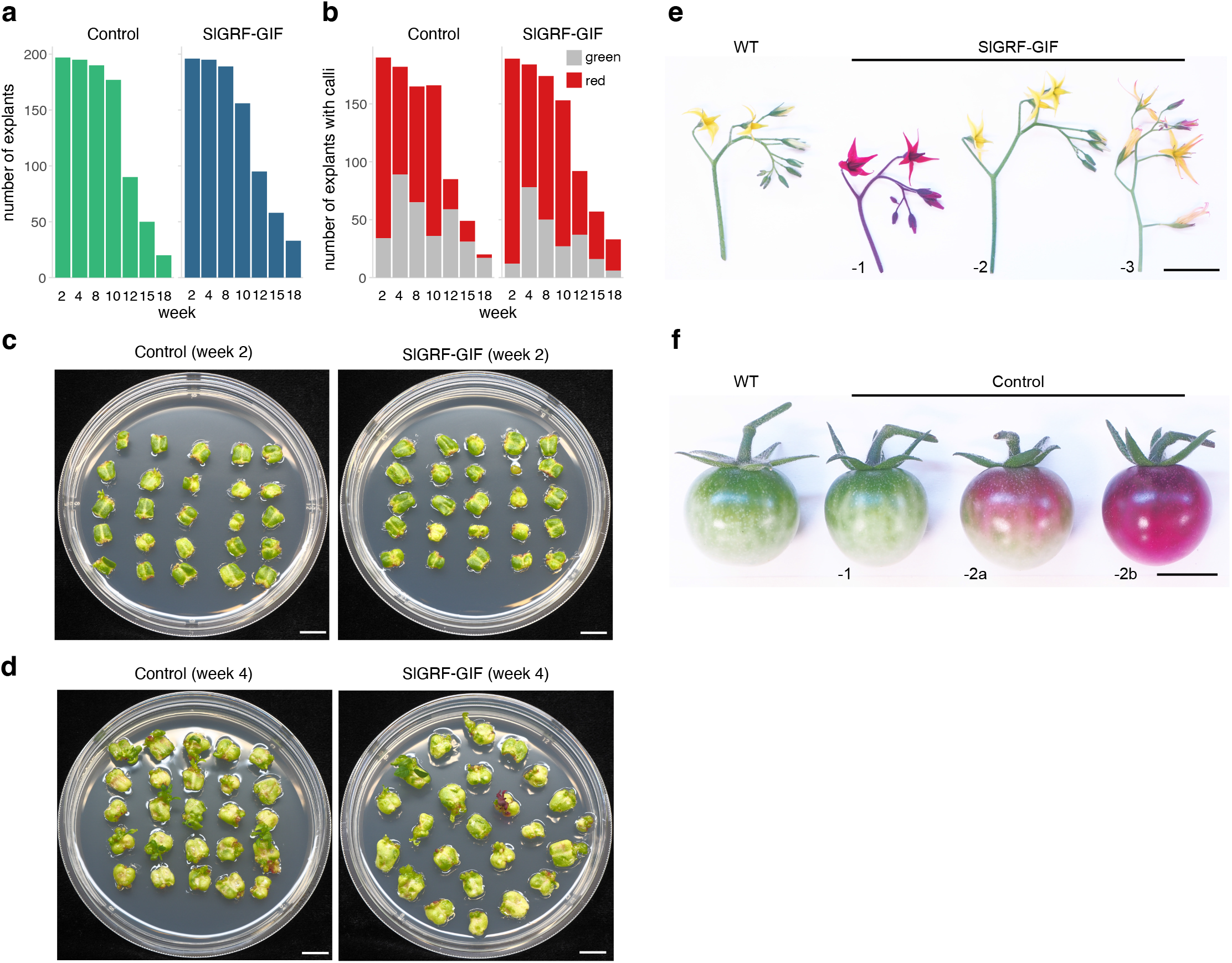
Transformation of the *RUBY* reporter with and without the *SlGRF-GIF* chimera. **a**, Quantification of the number of cotyledon explants on regeneration medium for the control and *SlGRF-GIF* construct. **b**, Quantification of explants with green and red calli on regeneration medium. **c-d**, Images of representative plates with explants at 2 weeks (c) and 4 weeks (d) after transformation. **e**, Images of representative inflorescences of wild type (WT) plants and *RUBY SlGRF-GIF* T0 individuals. **f**, Images of fruits at the mature green stage of WT plants and *RUBY* control T0 individuals. Scale bars in (c-f) represent 1 cm.

**Fig. S3:**
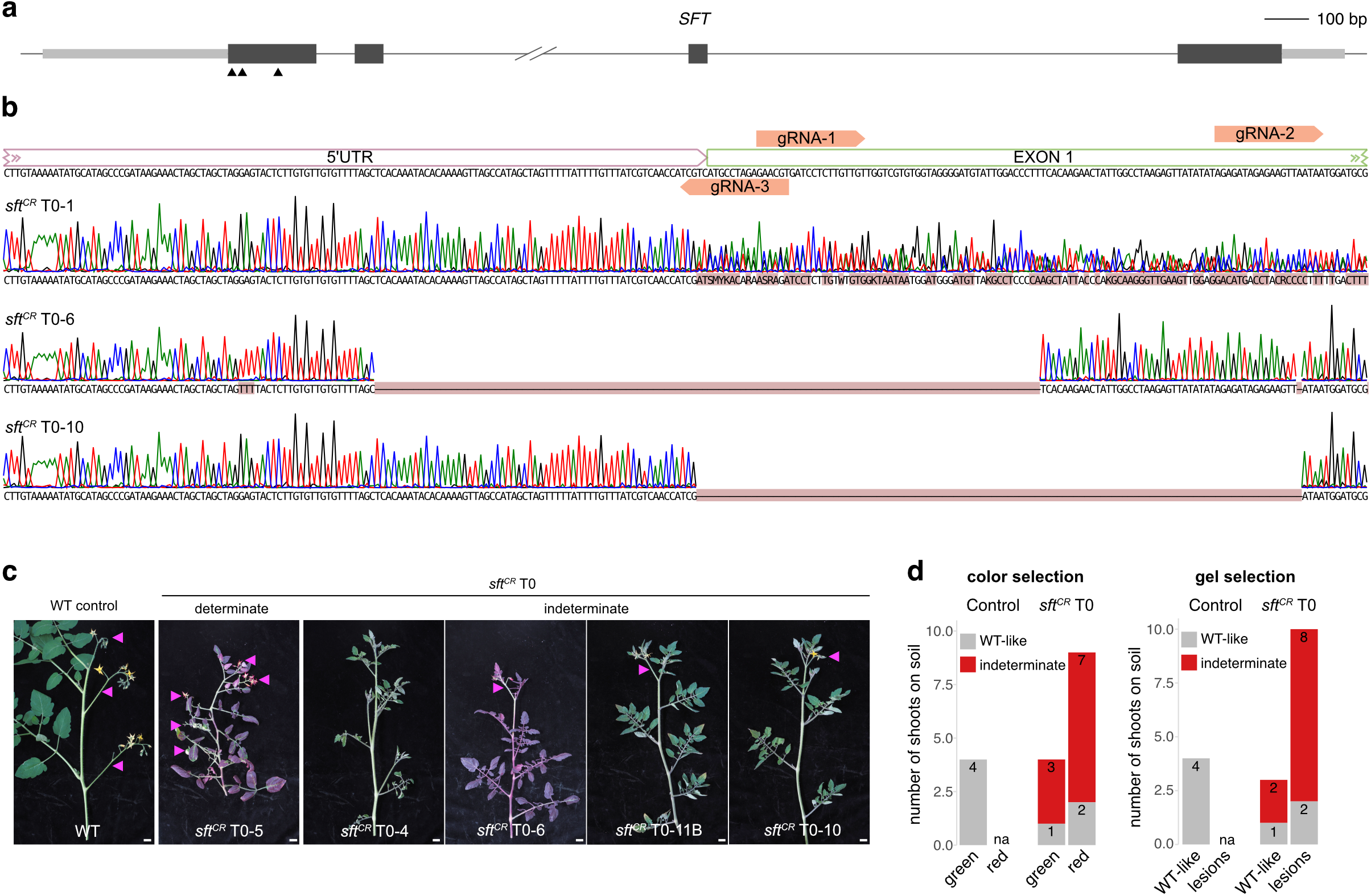
CRISPR-Cas targeting of *SFT* in combination with the morphogenic regulator *SlGRF-GIF* and the visible *RUBY* marker. **a**, Schematic representation of *SFT* with location of the Cas9 cleavage sites. Dark grey boxes represent exons and light grey boxes represent UTRs (untranslated regions). The Cas9 cleavage sites for guide RNAs are indicated with arrowheads. **b**, Verification of CRISPR-induced DNA lesions in *SFT* by Sanger sequencing. Mutated nucleotide positions are highlighted with red background. **c**, Images of shoots from the wild type (WT) and selected *sft^CR^* T0 individuals with determinate and indeterminate shoot growth. Scale bar represents 1 cm. **d**, Scoring determinate (WT-like) and indeterminate shoots on the WT control (control) plants and first-generation (T0) transgenic *sft^CR^* lines. Individuals were grouped by plant coloration (left) or DNA lesion detected by agarose gel (right). The number of individual plants is indicated in the bars. na, not applicable.

**Fig. S4:**
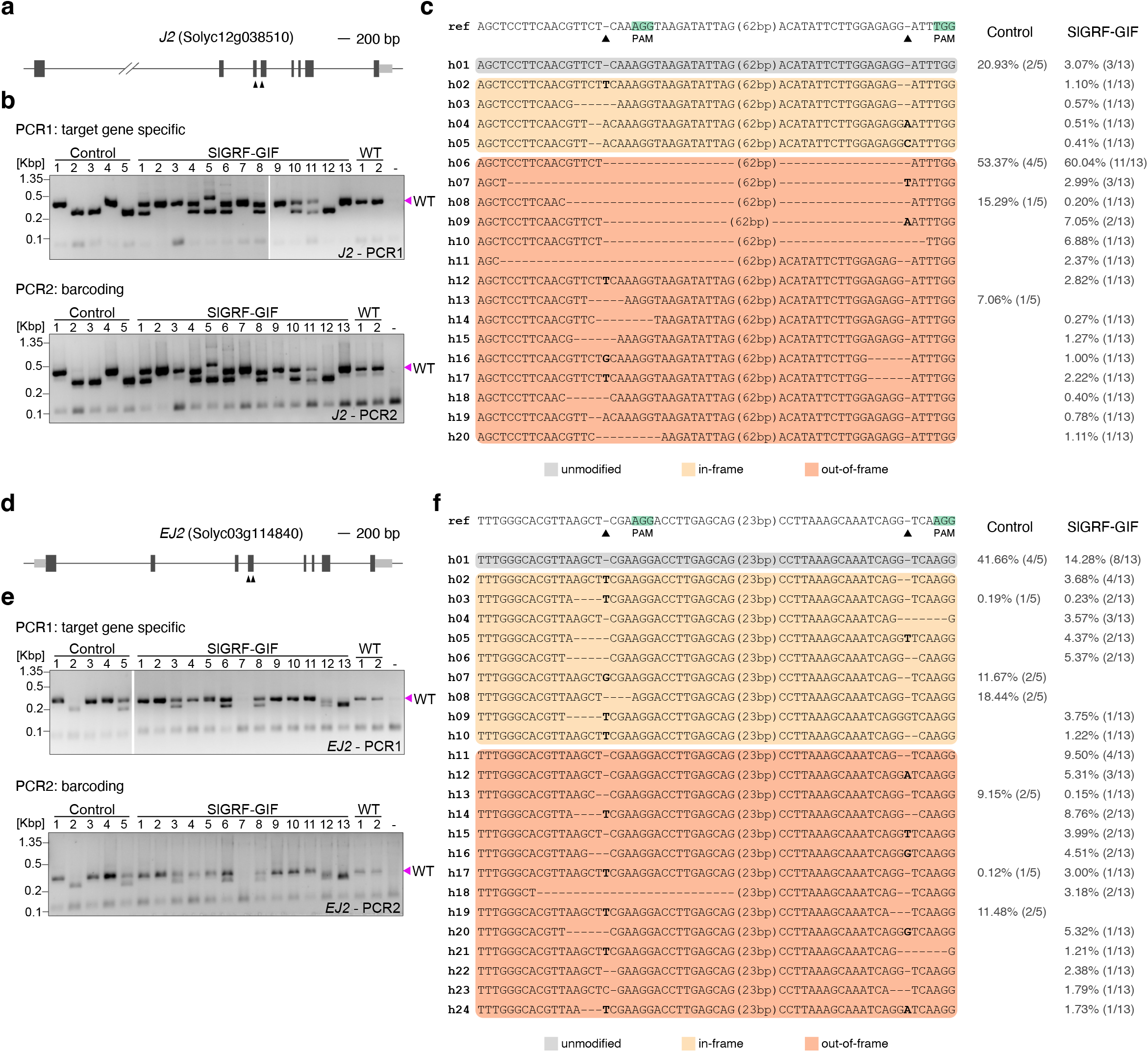
CRISPR-Cas multiplex targeting of *J2* and *EJ2* with and without the morphogenic regulator *SlGRF-GIF*. **a**, Schematic representation of *J2*. Dark grey boxes represent exons and light grey boxes represent UTRs (untranslated regions). The Cas9 cleavage sites for guide RNAs are indicated with arrowheads. **b**, Agarose gels showing target gene specific amplicons (top) and barcoded amplicons (bottom) of *J2* in the control and *SlGRF-GIF* regenerants with wild type (WT) and no DNA (-) controls. The size of the WT band is marked with a magenta arrowhead. Kbp, kilo base pairs. **c**, Haplotype sequences detected by SMAP- haplotype window analysis. The Cas9 cleavage sites for guide RNAs are indicated with arrowheads and PAMs are marked in green. Inserted bases are shown in bold, deleted bases are replaced by a dash, and sequence gap length is shown between parentheses. Haplotype frequency within the control and *SlGRF-GIF* experiments are shown together with the number of T0 transgenics in which the haplotype was detected between parentheses. ref, reference sequence. h, haplotype. PAM, protospacer adjacent motif. **d-f**, Schematic representation (**d**), agarose gels (**e**), and haplotype sequences (**f**) for *EJ2* as in (**a-c**).

**Fig. S5:**
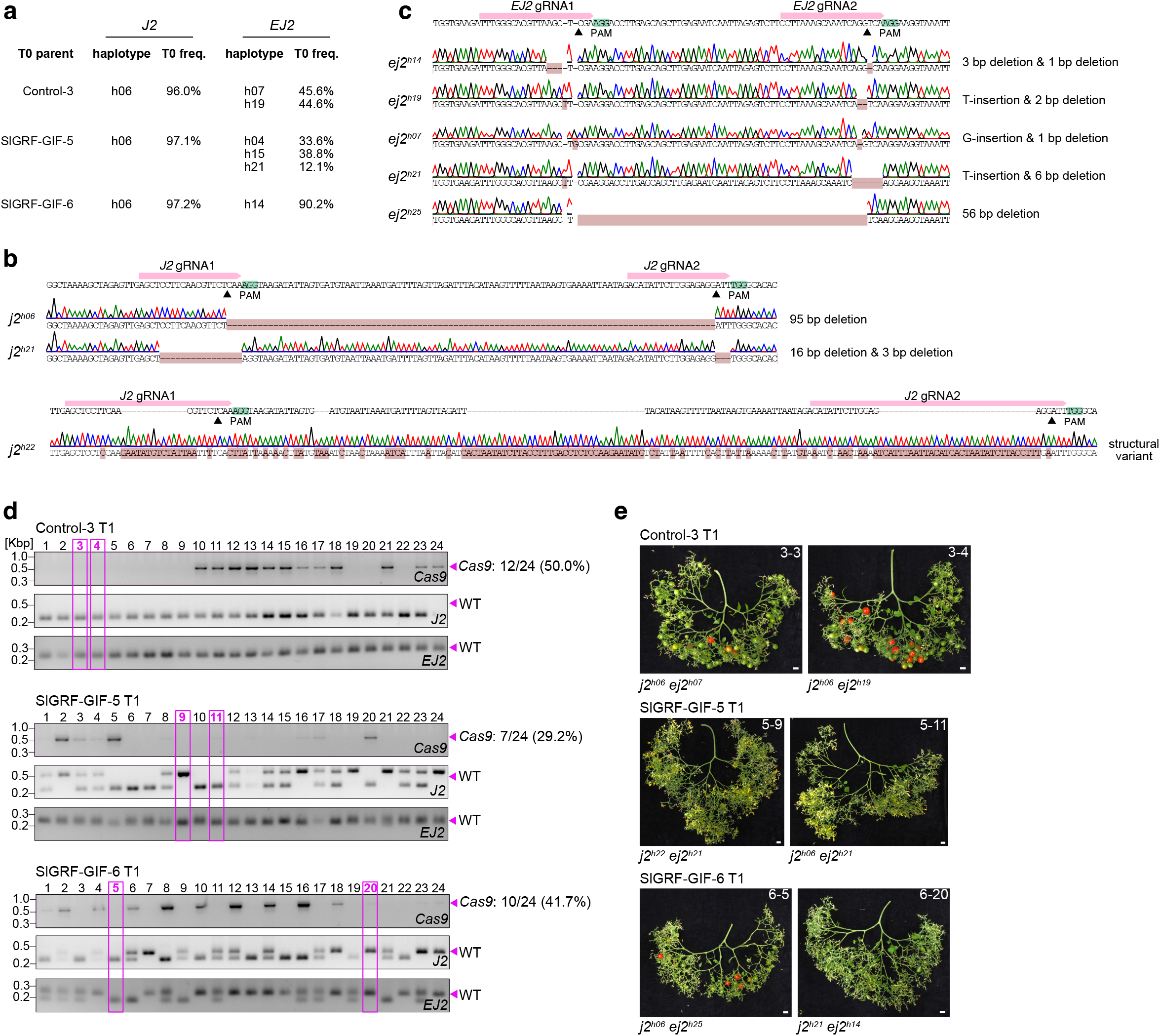
Recovery of *J2* and *EJ2* mutant haplotypes in non-transgenic T1 generation plants. **a**, Mutant haplotypes for *J2* and *EJ2* detected at frequencies > 5% in selected control and *SlGRF-GIF* T0 individuals. **b**, Verification of transmitted *j2* haplotypes by Sanger sequencing. Mutated nucleotide positions are highlighted with red background. The Cas9 cleavage sites for guide RNAs are indicated with arrowheads and PAMs are marked in green. PAM, protospacer adjacent motif. **c**, Verification of transmitted *ej2* mutant haplotypes as in (b). **d**, Agarose gels showing amplicons for the T-DNA (top), *J2* (middle), and *EJ2* (bottom) in control and *SlGRF-GIF* T1 families. The size of the expected T-DNA and WT amplicons are marked with magenta arrowheads. The number and percentage of Cas9-positive plants is indicated. Magenta boxes mark plants that were Sanger-sequenced and are shown in (e). Kbp, kilo base pairs. **e**, Images of detached inflorescences from two non-transgenic individuals per selected T1 family. Scale bars represent 1cm.

**Fig. S6:**
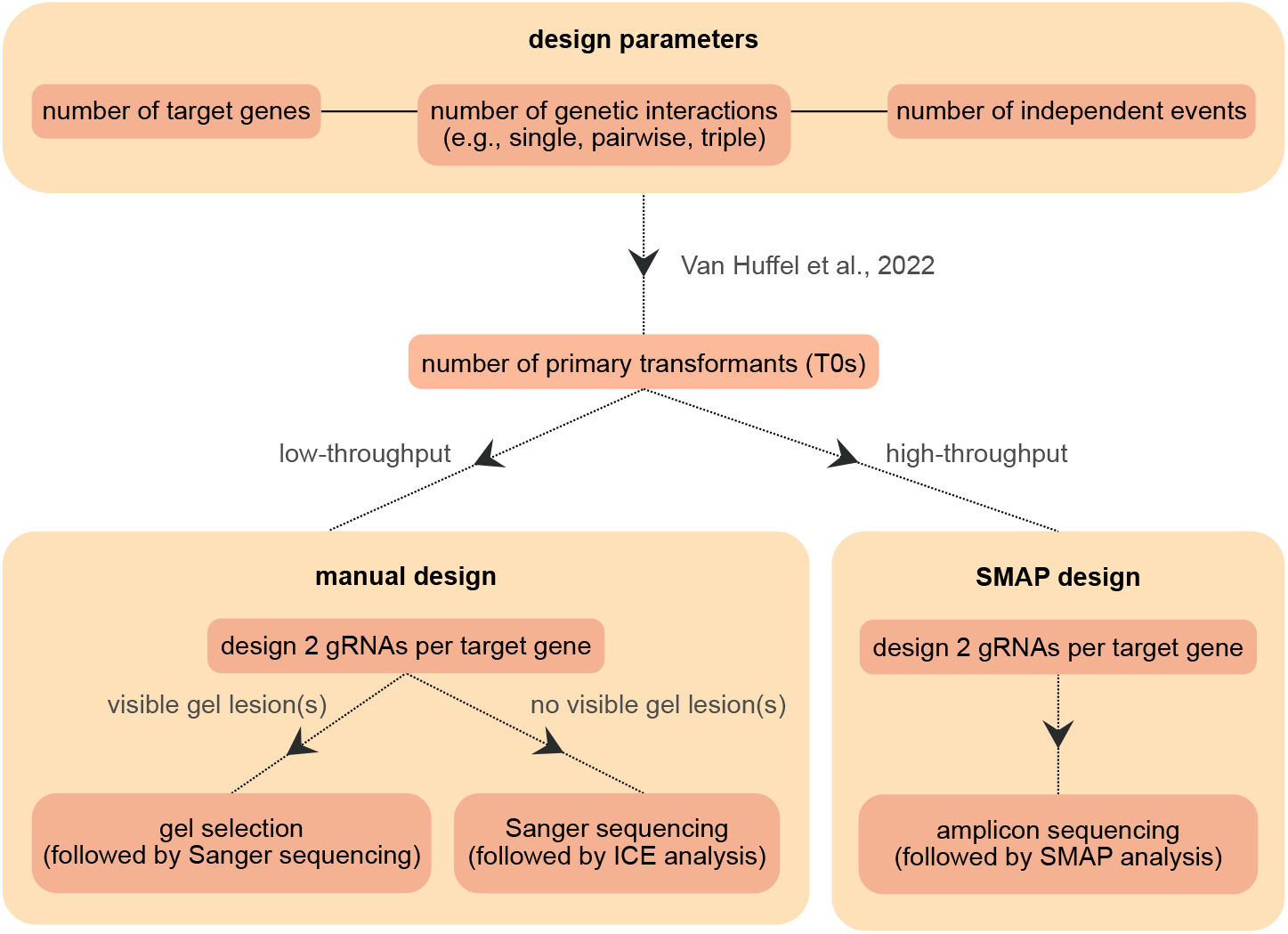
A flowchart for designing CRISPR-Cas experiments in tomato: Integrating T0 calculation, gRNA design, and genotyping strategy for generating gene loss-of-function lines. The target number of primary transformants (T0s) to be produced in a CRISPR-Cas experiment is determined by several factors, including the number of target genes, the number of genetic interactions to be investigated (single or higher-order mutants), and the number of desired independent genome editing events. A framework for calculating the number of T0s required based on these parameters is outlined in Van Huffel et al., 2022. The required number of T0 plants will inform the genotyping approach, and thereby also the gRNA design. For low-throughput experiments, such as the generation of single loss-of-function mutants, the gRNAs and genotyping primers can be designed manually to allow the induction and detection of large deletions that can be visualized on an agarose gel (followed by haplotype verification using Sanger sequencing). If no gel lesions are observed, amplicons containing the target region can be Sanger sequenced with a primer located ∼150-250 bp from the Cas cleavage site and quantitative sequence trace data can be decomposed using the Inference of CRISPR Editing (ICE) CRISPR Analysis Tool. High-throughput experiments, such as multiplex editing, can benefit from SMAP design (Develtere et al., 2023), which combines gRNA and genotyping primer design and offers optimal compatibility with amplicon sequencing. Haplotype sequences and their respective frequencies can be computed using SMAP-haplotype window analysis (Schaumont et al., 2022).

